# Skull bone marrow-derived immune cells infiltrate the damaged cortex and exhibit anti-inflammatory properties

**DOI:** 10.1101/2024.06.21.597827

**Authors:** Eman Soliman, Erwin Kristobal Gudenschwager Basso, Jing Ju, Andrew Willison, Michelle H. Theus

**Affiliations:** Department of Biomedical Sciences and Pathobiology, Virginia Tech, Blacksburg, VA 24061, USA; School of Neuroscience, Virginia Tech, Blacksburg, VA 24061, USA; Department of Physiology and Aging, COM, University of Florida, Gainesville, FL, 32610, USA; Translational Biology Medicine and Health Graduate Program, Virginia Tech, Blacksburg, VA, 24061, USA; Center for Engineered Health, Virginia Tech, Blacksburg, VA, 24061, USA

**Keywords:** Calvarium bone, Skull-derived monocytes, Blood-derived monocytes, controlled cortical impact, Innate immune response, Neuroinflammation

## Abstract

Identifying the origins and contributions of different immune cell populations following brain injury is crucial for understanding their roles in inflammation and tissue repair. This study investigated the infiltration and phenotypic characteristics of skull bone marrow-derived immune cells in the murine brain after TBI. We performed calvarium transplantation from GFP donor mice and subjected the recipients to controlled cortical impact (CCI) injury 14 days post-transplant. Confocal imaging at 3 days post-CCI revealed GFP+ calvarium-derived cells infiltrating the ipsilateral core lesional area, expressing CD45 and CD11b immune markers. These cells included neutrophil (Ly6G+) and monocyte (Ccr2+) identities. Calvarium-derived GFP+/Iba1+ monocyte/macrophages expressed the efferocytosis receptor MerTK and displayed engulfment of NeuN+ and caspase 3+ apoptotic cells. Phenotypic analysis showed that greater calvarium-derived monocyte/macrophages disproportionately express the anti-inflammatory arginase-1 marker than pro-inflammatory CD86. To differentiate the responses of blood- and calvarium-derived macrophages, we transplanted GFP calvarium skull bone into tdTomato bone marrow chimeric mice, then performed CCI injury 14 days post-transplant. Calvarium-derived GFP+ cells predominantly infiltrated the lesion boundary, while blood-derived TdTomato+ cells dispersed throughout the lesion and peri-lesion. Compared to calvarium-derived cells, more blood-derived cells expressed pro-inflammatory CD86 and displayed altered 3D morphologic traits. These findings uniquely demonstrate that skull bone-derived immune cells infiltrate the brain after injury and contribute to the neuroinflammatory milieu, representing a novel immune cell source that may be further investigated for their causal role in functional outcomes.

## Introduction

Traumatic brain injury (TBI) is a highly complex and devastating condition that places a significant burden on healthcare systems worldwide [1]. It is one of the primary causes of acute disability and death in the United States [2]. At the core of TBI’s pathology is neuroinflammation, which initiates a complex series of biochemical and cellular processes [3]. These processes extend beyond the initial trauma, causing secondary brain injury. Once the initial impact occurs, the brain tissue and blood-brain barrier become disrupted, triggering an inflammatory response. This response is characterized by the influx of immune cells and activation of resident microglia. The neuroinflammatory response is a crucial factor that can contribute to further damage and hinder recovery [3, 4]. Therefore, understanding the cellular response following trauma is vital in mitigating the secondary inflammatory response and improving the clinical outcomes of TBI.

Peripheral-derived monocytes are central to the inflammatory response following trauma, as they are recruited to the injured brain and differentiate into macrophages. Once matured, these macrophages engage in a range of activities mediating both protection and neuronal dysfunction. The neuroprotective or pro-resolving function of these phagocytes may be attributed to their ability to clear apoptotic cell debris formed following tissue damage via a process called efferocytosis [5, 6]. The detrimental effects, on the other hand, have been attributed to the production of pro-inflammatory and cytotoxic mediators [4]. It’s worth noting that these macrophages may originate from various peripheral sources, including border-associated regions and peripheral organs such as the spleen and bloodstream, each potentially contributing differently to the inflammation and repair processes.

The bone marrow niches located in close proximity to the brain may also serve as reservoirs for myeloid cells that support the meninges and CNS parenchyma [7]. These cells are distinct from blood-derived myeloid cells, especially in the context of neuroinflammation, as they are supplied by the adjacent bone marrow and exhibit unique signatures [7, 8]. Their strategic placement at the CNS borders positions them ideally for immediate immune responses. Skull bone marrow myeloid cells communicate directly with the glymphatic system of CNS through specialized channels and are believed to actively participate in immune surveillance and neuroimmune communication [9]. However, it is yet to be determined how skull bone marrow-derived cells respond to brain trauma. Given their proximity to the dura, they may infiltrate damaged tissue following injuries and elicit a unique inflammatory response from their blood-derived counterparts [8]. The current study is the first of its kind to characterize skull bone marrow-derived myeloid cell infiltration in the brain after traumatic injury and to distinguish them from those derived from the circulation.

## Methods

### Animals

Male Rosa26-tdTomato and GFP mice were purchased from Jackson Laboratory, Bar Harbor, ME, and backcrossed 10 generations and maintained on the CD1 background. All mice were maintained in groups of four to five per cage at a facility accredited by AAALAC. They were kept on a 12-hour light-dark cycle, with unrestricted access to food and water. Tail snips were utilized for genotyping as previously outlined [10, 11]The study protocols were approved by the Virginia Tech Institutional Animal Care and Use Committee (IACUC; #21-044).

### Chimeric bone marrow transplantation

Generation of GFP or tdTomato bone marrow chimeric mice was carried out as previously described [12, 13]. Male CD1 recipient mice were anesthetized by subcutaneous injection of ketamine (100 mg/kg) and xylazine (10 mg/kg) and then subjected to two sessions of X-ray irradiation, each delivering 500 rad and spaced six hours apart (Radsource, Brentwood, TN), with head shields. Twenty-four hours after irradiation, these mice were intravenously injected with 1-5 million bone marrow cells harvested from age-matched male Rosa26*-*tdTomato or *-*GFP donor mice [14]. Mice received water mixed with gentamycin (1 mg/ml) for three days before irradiation and two weeks following the bone marrow transplantation.

### Calvarium bone-flap transplantation

Calvarium transplantation was performed as described [7]. Briefly, recipient mice (CD1 or tdTomato chimeric at 30 days post bone marrow transplant) were anesthetized by subcutaneous injection of ketamine (100 mg/kg) and xylazine (10 mg/kg), secured in a stereotaxic frame, and kept on a heating pad set at 37°C to maintain body temperature throughout the surgery. The procedure involved shaving the heads of recipient mice, making a midline incision in the skin, and exposing the skull. A cranial window (4 mm by 6 mm) was made on the parietal and interparietal bones using a portable electric drill, with intermittent cooling using saline. The bones were carefully removed without damaging the underlying dura. Simultaneously, age-matched GFP mice were euthanized to harvest calvarium bone flaps for transplantation. The donor skull pieces containing the bone marrow were collected, connective tissues were kept intact to enhance graft survival, and meninges were removed. The bones were then positioned and secured within the host’s surgical window using cyanoacrylate glue. The skin was sutured, and the mice were kept warm until recovery.

### Controlled cortical impact (CCI) injury

At 14 days post-calvarium transplantation, CCI was carried out as previously described [6, 15-17]. Briefly, mice were anesthetized using a subcutaneous injection of ketamine (100 mg/kg) and xylazine (10 mg/kg) and then positioned in a stereotaxic frame. After preparing the scalp by shaving and sanitizing, a midline incision was made to expose the skull, where a 4 mm craniectomy was performed over the right parietal-temporal cortex (− 2.5 mm A/P and 2.0 mm lateral from the bregma) with a portable drill. The injury was induced using an eCCI-6.3 device (Custom Design & Fabrication, LLC) with parameters of 5.0 m/s velocity, 2 mm depth, and 150 ms duration. The incision site was sutured, and mice were monitored until they fully recovered from anesthesia.

### Cardiac perfusion, fixation, and brain serial coronal sectioning

At the end of the experiment, mice were administered 2000 units/kg of heparin and subsequently euthanized with lethal doses of ketamine (150 mg/kg) and xylazine (15 mg/kg) cocktail. Cardiac perfusion was performed at a 5 ml/min flow rate using a Gilson MiniPuls3 peristaltic pump (Gilson Scientific, Bedfordshire, UK). Initially, 10 ml of heparin solution (20 units/ml in 1X PBS) was perfused through the left ventricle to clear the blood, then followed by 50 ml of ice-cold 4% paraformaldehyde (PFA) in PBS (pH 7.5). The brains were then harvested and cryopreserved using a sucrose gradient before being snap-frozen and embedded in Tissue Tek O.C.T (Sakura, Torrance, CA) and stored at - 80°C. Serial coronal brain sections (− 1.1 to − 2.6 mm A/P) were collected using a CryoStar NX70 cryostat (Thermo Fisher Scientific, USA). Five 30μm thick tissue sections spaced 450μm apart were mounted on charged slides, heat dried, and then stored at -80°C.

### Immunohistochemical analysis and cell counts

Serial coronal sections were blocked using 2% cold water fish skin gelatin (Sigma, Inc. St. Louis, MO) and then incubated with primary antibody solutions (1:100) made in the blocking solution. The sections were washed with 1X PBS and incubated with the corresponding secondary antibodies, washed then mounted using a medium containing DAPI counterstain (Southern Biotech, Birmingham, AL). The primary antibodies used include rabbit anti-CD45 (Cell Signaling, Danvers, MA, USA), rat anti-CD11b (Abcam, Waltham, MA, USA), rabbit anti-CCR2 (Abcam, Waltham, MA, USA), rat anti-CD86 (Invitrogen, Waltham, MA, USA), rabbit anti-Iba1(FUJIFILM Wako Chemicals U.S.A. Corporation), rat anti-Iba1(Abcam, Waltham, MA, USA), chicken anti-Arg1 (Phosphosolutions, Aurora, CO, USA), rat anti-Ly6G (Biolegend, San Diego, CA, USA), rabbit anti-MERTK (Abcam, Waltham, MA, USA), rat anti-NeuN (Abcam, Waltham, MA, USA), and rabbit anti-active caspase 3 (R&D systems, Minneapolis, MN, USA). The secondary antibodies used include donkey anti-rabbit 647, donkey anti-rabbit 594, and donkey anti-rat 647, donkey anti-rat 594, donkey anti-chicken 647, donkey anti-chicken 594 (all from Thermofisher, USA). Images were captured using a Zeiss 880 confocal microscope (Carl-Zeiss, Oberkochen, Germany). Confocal z-stack images were imported into IMARIS software (Bitplane, Zurich, Switzerland) for 3D reconstruction and quantification of cell morphology. Surface rendering of peripheral-derived macrophages was performed with a smoothing factor of 0.1 μm. An absolute thresholding method was applied in a blinded fashion, and surfaces smaller than 10 μm^3^ were excluded. Cell counting was blindly assessed using image J (2.9.0 Fiji software) or the optical fractionator probe function of Stereoinvestigator (MicroBrightField, Williston, VT, USA) as previously described [15, 16].

### Statistical Analysis

Data are expressed as mean ± standard error of the mean (SEM). Statistical analysis was performed using Graph Pad Prism software version 10 (La Jolla, CA, USA). To analyze differences between the groups, we used the student’s unpaired two-tailed t-test for comparing two experimental groups and one-way analysis of variance (ANOVA) followed by Bonferroni’s post hoc test for comparing more than two groups. P<0.05 was considered statistically significant.

## Results

### Calvarium-derived immune cells infiltrate the brain following CCI injury

Identifying the origin and the specific contributions of different immune cell populations following TBI is essential for understanding their roles in inflammation and the healing process. In this study, we investigated the infiltration of immune cells derived from the skull bone marrow into the brain after TBI by performing a series of experiments involving Calvarium bone-flap transplantation followed by CCI injury. Calvarium bone was isolated from GFP donor mice and transplanted into CD1 recipient mice 14 days before the induction of CCI injury (Fig. 1A). Specifically, a 4 mm by 6 mm section of calvarium bone was harvested from GFP donor mice and positioned into age-matched CD1 recipient mice with calvarium windows located at the parietal and interparietal bones. Fourteen days post-transplant, an examination of the CCI craniectomy site revealed a healed bone structure with both GFP+ and GFP-regions, indicating successful transplantation (Fig. 1B).

**Figure 1:**
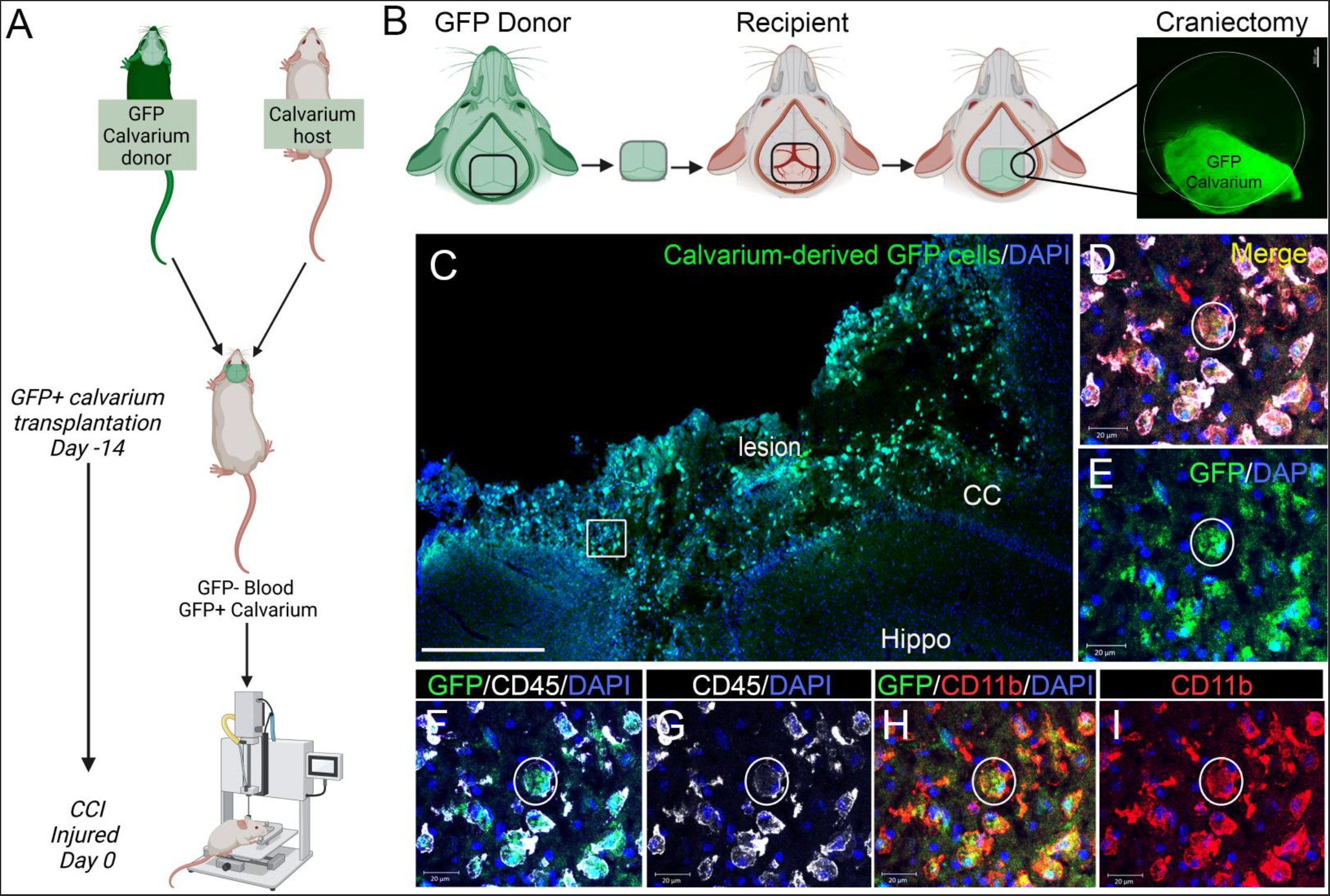
Immune cells from skull bone marrow infiltrate the brain after CCI injury. **(A)** Experiment outline for calvarium bone-flap transplantation and CCI injury. Calvarium was isolated from GFP donor mice and transplanted into CD1 recipient mice 14 days before the induction of CCI injury. **(B)** A 4 mm by 6 mm section of calvarium bone was isolated from a GFP donor mouse and transplanted into an age-matched CD1 recipient mouse with a calvarium window created in the parietal and interparietal bones. Fourteen days post-transplant, CCI craniectomy revealed a healed bone structure containing both GFP+ and GFP-regions. **(C)** Representative confocal image showing calvarium-derived GFP+ cells (green) infiltrating the injured tissue (lesion) and the corpus callosum (CC) of the ipsilateral cortex at 3 days post-CCI injury (5X magnification). **(D-I)** Representative images showing calvarium-derived GFP+ cells (green) expressing CD45 (white) and CD11b (red).

At 3 days post-CCI injury, calvarium-derived GFP+ cells infiltrated the ipsilateral cortex and were detected at both the lesion core, boundary, and the corpus callosum (CC) (Fig. 1C). Further immunohistochemical analysis revealed that these GFP+ cells expressed CD45 and CD11b markers, confirming their identity as immune cells (Fig. 1D-1I). Our findings highlight the presence of calvarium-derived immune cells in the early response to brain injury, which may contribute to the cellular landscape within the injured cortex.

### Calvarium-derived immune cells comprise neutrophil and monocyte populations

Previous studies have reported that the skull and vertebral bone marrow supply the mouse meninges with a pool of monocytes and neutrophils [7]. To determine if skull bone marrow-derived immune cells infiltrating the injured cortex contain monocyte and neutrophil populations, we stained the coronal sections of mice that underwent GFP calvarium transplantation followed by CCI injury with a monocyte marker, Ccr2, and a neutrophil marker, Ly6G (Fig. 2). Confocal image analysis revealed the presence of GFP+CD45+Ly6G+ calvarium-derived neutrophils (Fig. 2A) and GFP+CD11b+Ccr2+ calvarium-derived monocytes (Fig. 2B) in the ipsilateral cortex at 3 days post-CCI injury. Additionally, GFP-CD11b+Ccr2+ non-skull-derived monocytes were also observed in the injured cortex (Fig. 2B), highlighting the diverse origins of cells participating in the immune response to injury.

**Figure 2:**
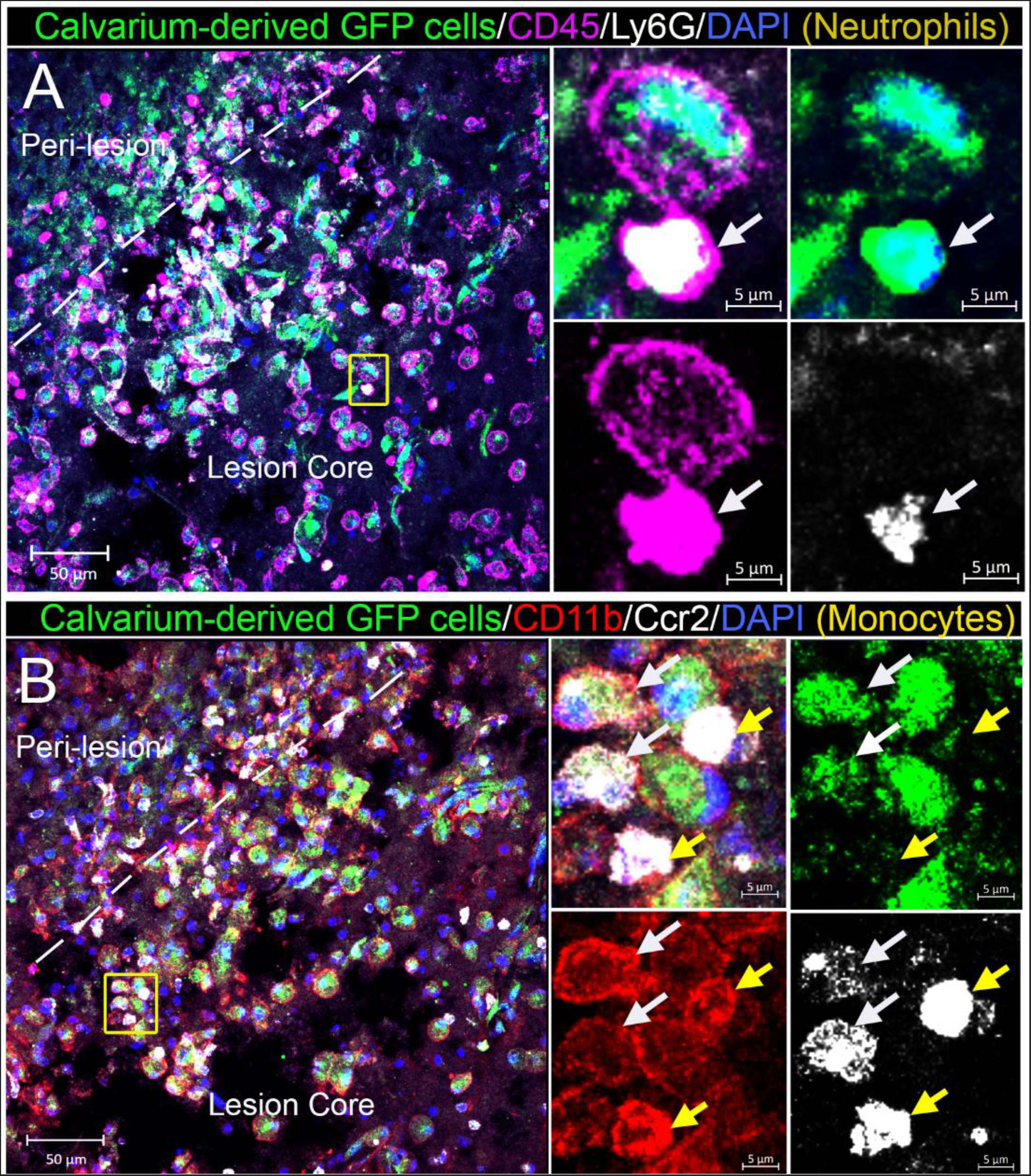
Immune cells originating from the skull bone marrow encompass both neutrophils and monocytes. Mice were subjected to GFP calvarium transplant followed by CCI injury. **(A)** Representative confocal image and region of interest (square) showing calvarium-derived neutrophils (white arrow) expressing GFP (green), CD45 (purple), and Ly6G (white) infiltrating the ipsilateral cortex at 3 days post-CCI injury. **(B)** Representative confocal image showing two populations of CD11b+/Ccr2+ monocytes infiltrating the ipsilateral cortex at 3 days post-CCI injury. The region of interest shows calvarium-derived monocytes (white arrow) express GFP+ (green), CD11b (red), and Ccr2 (white), and non-skull-derived monocytes (yellow arrow) express CD11b (red) and Ccr2 (white) without GFP.

### Calvarium-derived macrophages participate in cell debris clearance

Our previous studies have demonstrated that peripheral-derived macrophages express the efferocytosis receptor, MerTK, and participate in the clearance of apoptotic neurons in the brain following injury [18]. However, these studies did not distinguish between blood-derived and skull-bone marrow-derived macrophages. To further investigate whether skull-bone marrow-derived macrophages contribute to efferocytosis following TBI, we used the coronal sections of injured mice that underwent GFP calvarium bone-flap transplantation and detected the engulfment of neurons (NeuN-positive) and apoptotic cells (activated caspase 3-positive) in the injured cortex. We observed that GFP+IBA1+ calvarium-derived macrophages express MerTK in the ipsilateral cortex 3 days post-CCI injury (Fig. 3A-G). In addition, MerTK-expressing GFP+ cells engulfed NeuN+ neurons (red) at the lesion core (Fig. 3H; yellow arrow). Additionally, IBA1+GFP+ macrophages engulfed caspase 3+ (Casp3, white) cells, indicating their involvement in clearing apoptotic cells at the lesion core (Fig. 3I; cyan arrow). These findings suggest that calvarium-derived macrophages may contribute to post-injury tissue remodeling by participating in the clearance of neuronal debris and apoptotic cells in the injured brain.

**Figure 3:**
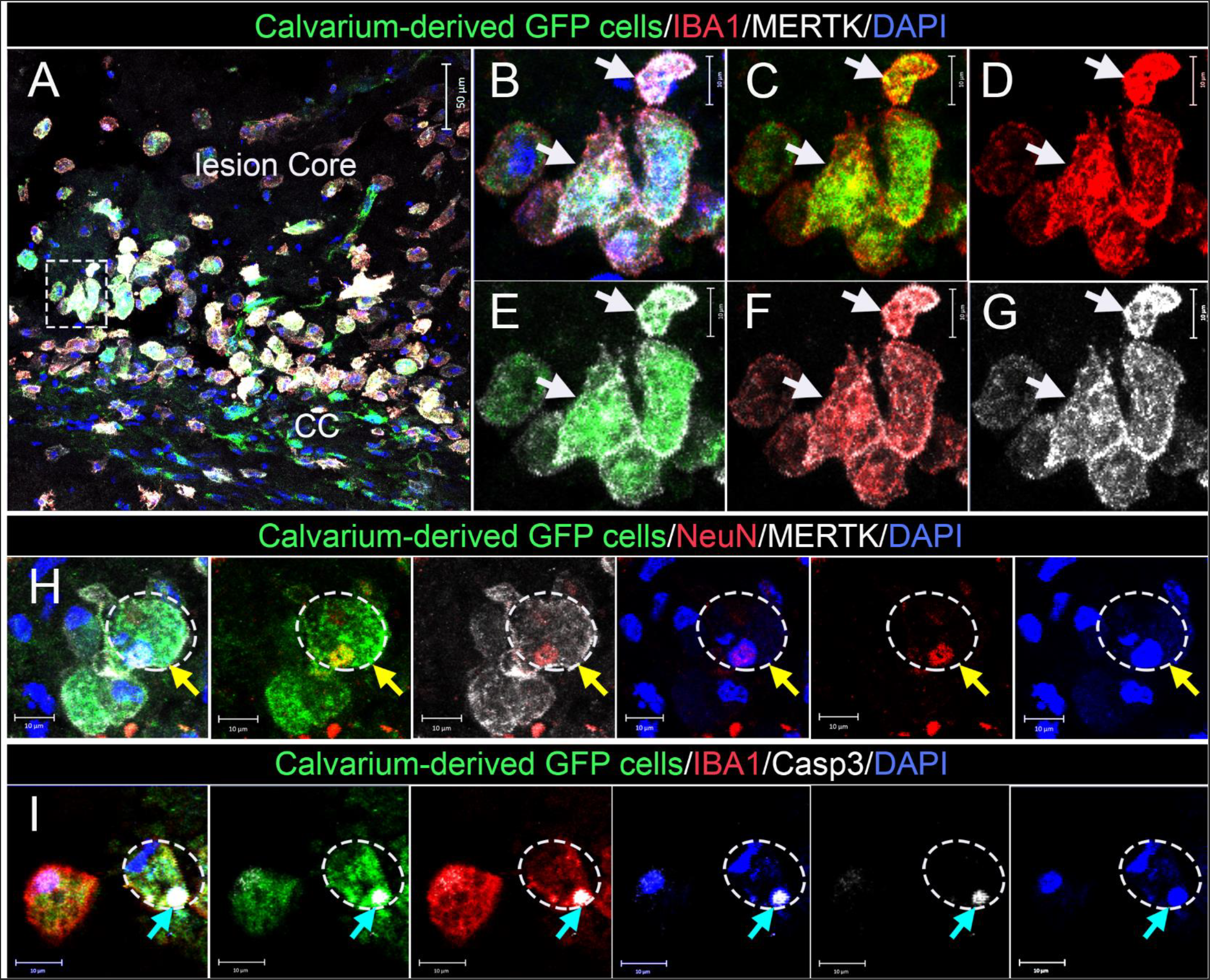
Skull-bone marrow-derived macrophages contribute to the clearance of neuronal debris in the injured brain. Mice were subjected to GFP calvarium bone-flap transplantation followed by CCI injury. **(A-G)** Representative confocal image and region of interest (square) showing calvarium-derived macrophages (white arrows) expressing GFP (green), IBA1 (red), and efferocytosis receptor MerTK (white) in the ipsilateral cortex at 3 days post-CCI injury. **(H)** Representative confocal images showing MerTK+ (white) GFP+ (green) calvarium-derived cells engulfing NeuN+ (red) neurons at the lesion core (yellow arrow). **(I)** Representative confocal images showing IBA1+ (red) GFP+ (green) calvarium-derived macrophages engulfing caspase 3+ (Casp3, white) cells at the lesion core (cyan arrow).

### Phenotypic state of infiltrating Calvarium-derived macrophages

Monocytes/macrophages can exhibit either pro-inflammatory or anti-inflammatory phenotypes after brain injury. The phenotypic response of calvarium-derived monocyte/macrophages was evaluated in the ipsilateral cortex at 3 days post-CCI injury. Immunohistochemical analysis revealed that calvarium-derived GFP+ monocyte/macrophages express CD86+ proinflammatory (Fig. 4A-4E) or Arg1+ anti-inflammatory (Fig. 4G-4K) markers. Non-biased stereological quantification shows that ∼64% of GFP+ calvarium-derived cells were monocyte/macrophages (GFP+IBA1+; Fig. 4F). Additionally, ∼65% of these cells express anti-inflammatory marker Arg1 (GFP+IBA1+Arg1+); while only ∼17% express pro-inflammatory CD86 marker (GFP+IBA1+CD86+, Fig. 4L).

**Figure 4:**
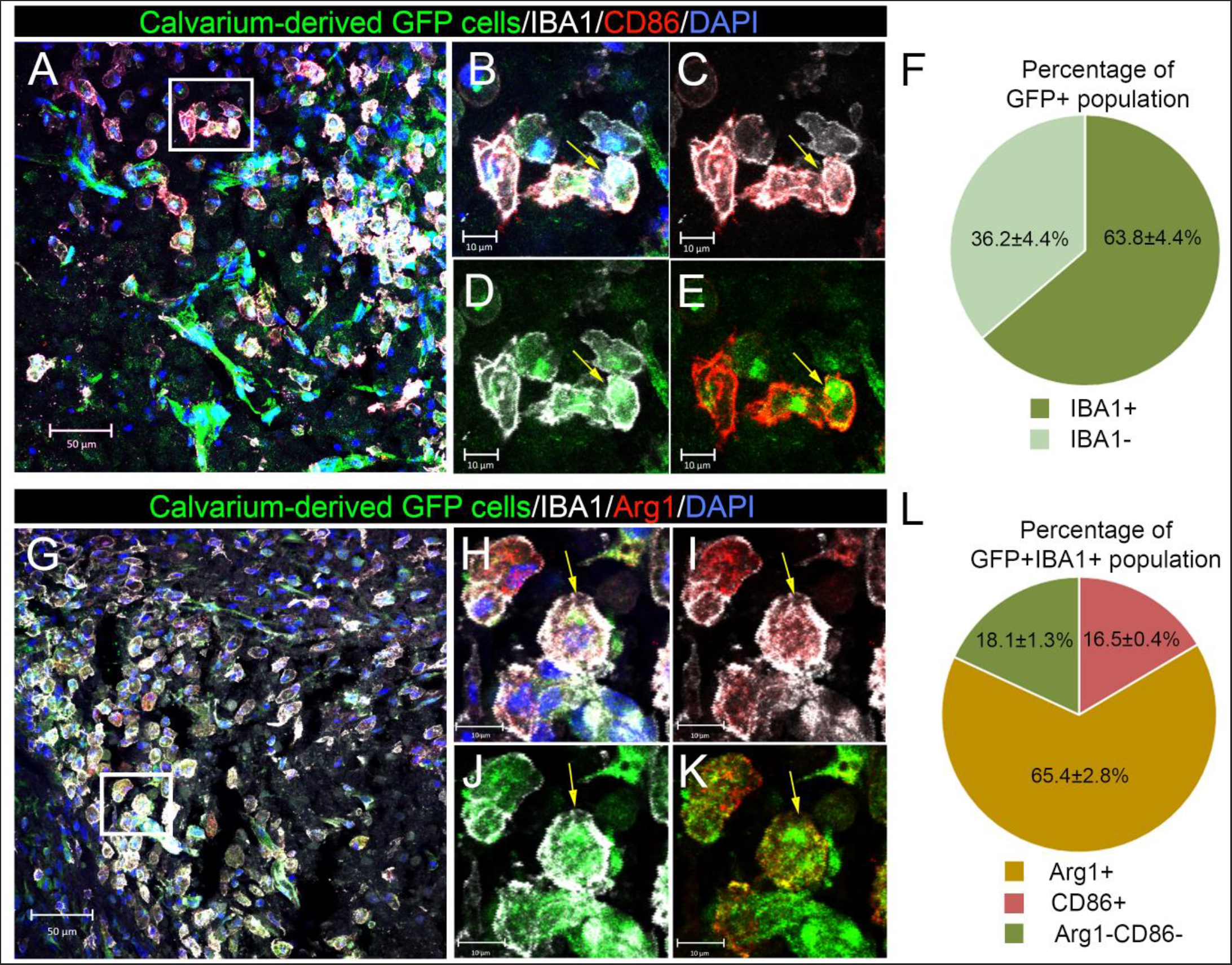
Phenotypic characterization of skull-bone marrow-derived macrophages infiltrating the injured brain. Mice were subjected to GFP calvarium bone-flap transplantation followed by CCI injury. **(A-E)** Representative confocal image and region of interest (square) showing calvarium-derived macrophages (yellow arrows) expressing GFP (green), IBA1 (white), and a pro-inflammatory phenotype marker, CD86 (red) in the ipsilateral cortex at 3 days post-CCI injury. **(F)** A pie chart showing the percentage of calvarium-derived macrophage (GFP+IBA1+) population within the total GFP+ cell population. **(G-K)** Representative confocal images and region of interest (square) showing calvarium-derived macrophages (yellow arrows) expressing GFP (green), IBA1 (white), and an anti-inflammatory phenotype marker, Arg1 (red) in the ipsilateral cortex at 3 days post-CCI injury. **(L)** A pie chart showing the percentage of calvarium-derived pro-inflammatory macrophages (GFP+IBA1+CD86+) and anti-inflammatory macrophages (GFP+IBA1+Arg1+) within the macrophage (GFP+IBA1+) population.

### Characterization of blood- and calvarium-derived monocyte/macrophages in the damaged cortical milieu

To characterize the histological nature of blood-versus calvarium-derived macrophages in the damaged cortex, we performed GFP calvarium bone-flap transplantation in tdTomato bone marrow chimeric mice at 30-day post-adoptive transfer and subsequent CCI injury at 14 days bone transplant (Fig. 5A). Confocal image analysis showed that calvarium-derived GFP+ cells predominantly infiltrated the lesion boundary at 3 days post-CCI injury, while blood-derived TdTomato+ cells were more dispersed in the core and peri-lesion area of the ipsilateral cortex (Fig. 5B-5D). The total number of blood-derived TdTomato+ cells infiltrating the ipsilateral cortex was significantly higher than calvarium-derived GFP+ cells (Fig. 5M). Further characterization of these cells revealed distinct phenotypic differences. A greater number of blood-derived TdTomato+ cells expressed pro-inflammatory CD86, while an increased number of calvarium-derived GFP+ cells expressed anti-inflammatory Arg1 (Fig. 5M). The percentage of Arg1-expressing cells was significantly higher among the GFP+ population, while the percentage of CD86-expressing cells was higher among the tdTomato+ population (Figure 5N).

**Figure 5:**
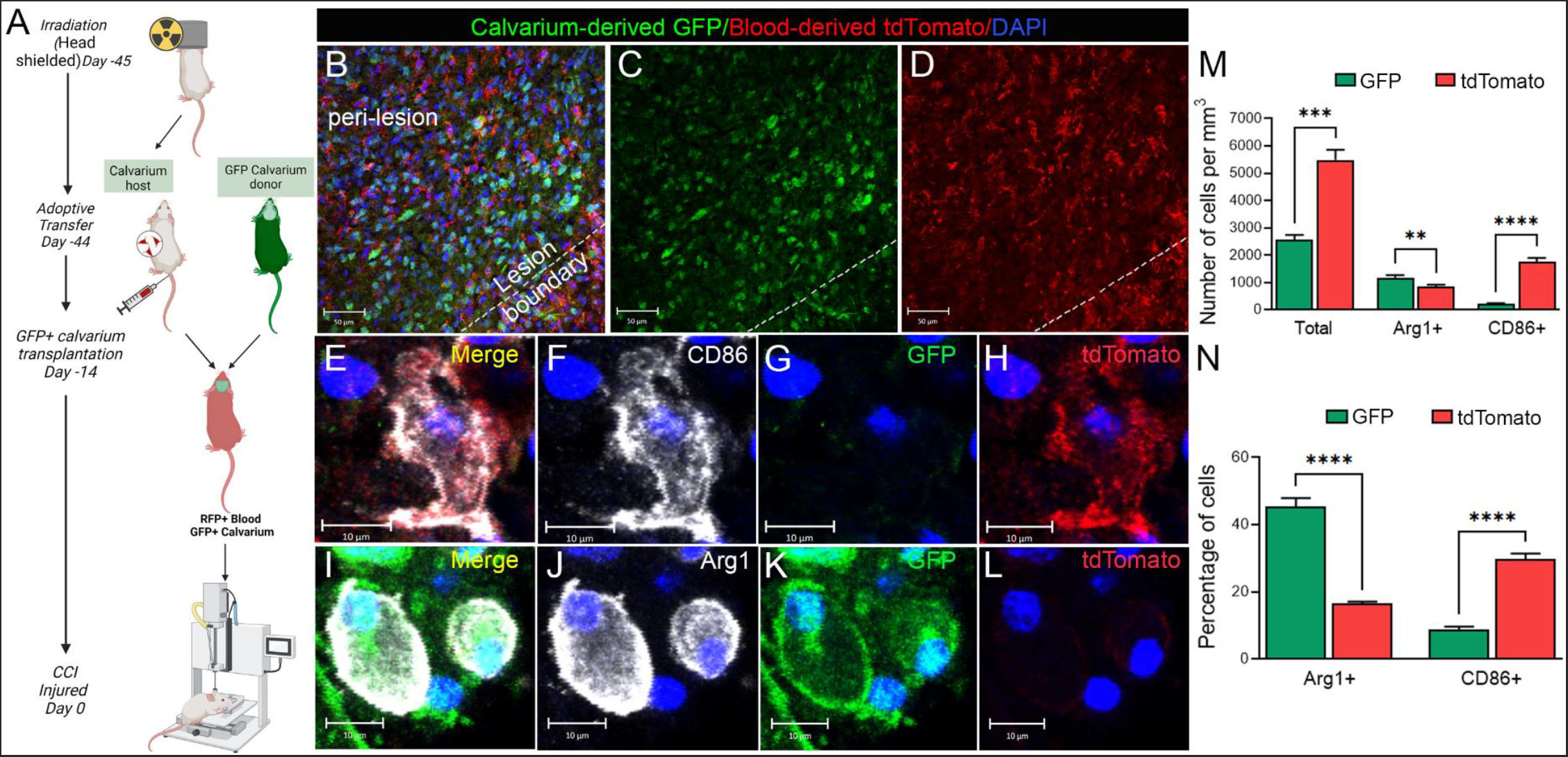
Differentiating the responses of blood- and skull-bone marrow-derived monocyte/macrophages in the brain following CCI injury. **(A)** Experiment outline for adoptive transfer, calvarium bone-flap transplantation, and CCI injury. CD1 wild-type mice were subjected to X-ray irradiation with head-shielding 45 days before CCI, then received TdTomato bone marrow cells 24 hours after irradiation to generate TdTomato adoptive transfer mice. Thirty days after the bone marrow transfer, TdTomato adoptive transfer mice were subjected to calvarium transplantation and received GFP calvarium from GFP donor mice. Two weeks later, mice with tdTomato+ (anti-RFP) blood and GFP+ calvarium were subjected to CCI injury. **(B-D)** Representative confocal images showing calvarium-derived GFP+ cells (green) and blood-derived TdTomato+ cells (red) infiltrating the lesion boundary and peri-lesion areas of the ipsilateral cortex at 3 days post-CCI injury. More GFP+ cells are observed at the lesion boundary, while TdTomato+ cells are more dispersed in the peri-lesion area. **(E-H)** Representative images showing blood-derived TdTomato+ cells (red) expressing CD86 (white). **(I-L)** Representative images showing calvarium-derived GFP+ cells (green) expressing Arg1 (white). **(M)** The total number of GFP+ or tdTomato+ cells expressing Arg1 or CD86 was counted in the ipsilateral cortex at 3dpi using the optical fractionator, Stereoinvestigator. **(N)** The percentage of Arg1- or CD86-expressing cells was calculated from the total number of GFP+ and RFP+ cells. n=4 mice/group. **P<0.01; ***P<0.001, ****P<0.0001. Multiple t-tests.

Interestingly, the morphology of CD86+ TdTomato+ cells was notably different from that of Arg1+GFP+ cells (Fig. 5E-5L). Therefore, morphological analysis of calvarium-and blood-derived cells was performed using IMARIS software, and 3D reconstructions of the lesion boundary and peri-lesion areas were conducted using z-stack confocal images for the ipsilateral cortex of injured mice at 3 days post-CCI injury. These reconstructions revealed distinct localization patterns: GFP+ cells were predominantly located at the lesion boundary, whereas TdTomato+ cells were more prevalent in the peri-lesion area (Fig. 6A and 6B). Quantification of the total volume of fluorescence indicated that GFP+ cells exhibited a significantly greater volume than tdTomato+ cells at the lesion boundary (p < 0.01) (Fig. 6C), suggesting that calvarium-derived cells occupy more space and potentially have different functional characteristics. Analysis of cell sphericity showed that GFP+ cells had higher sphericity values compared to TdTomato+ cells in both the lesion boundary and peri-lesion areas (p < 0.05) (Fig. 6D), indicating that calvarium-derived macrophages are more ameboid. Additionally, the average surface area of TdTomato+ cells was significantly greater than that of GFP+ cells in the peri-lesion area (p < 0.05) (Fig. 6E), indicating their processes may be more extensive. Taken together, the distinct spatial distribution, morphological characteristics, and phenotypic markers expressed by calvarium-versus blood-derived macrophages underscore the potentially diverse functional roles of these populations in the inflammatory response following brain injury.

**Figure 6:**
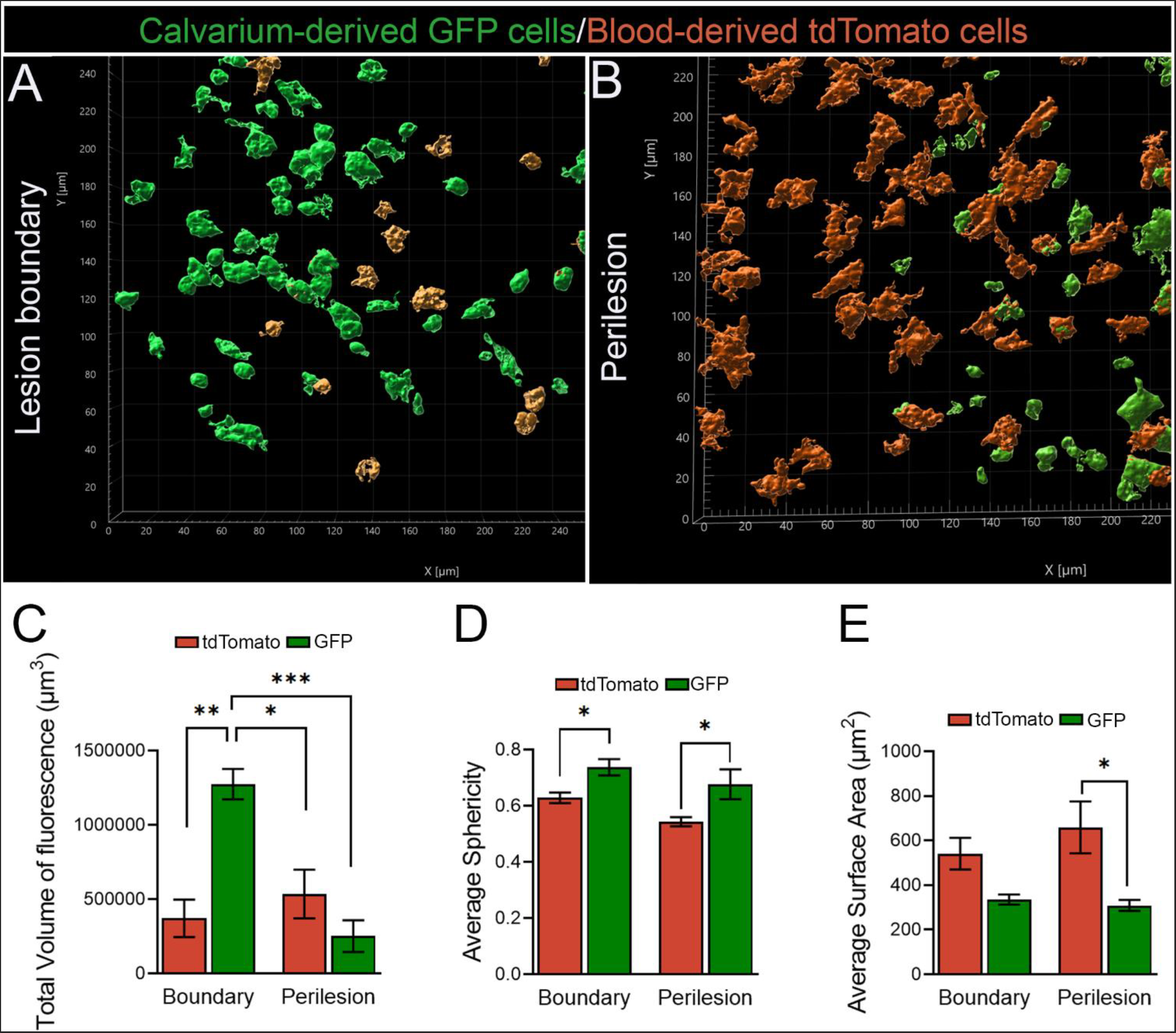
Morphological analysis of blood-derived TdTomato cells and skull-bone marrow-derived GFP cells in the injured brain following CCI. **(A, B)** 3D reconstructions of calvarium-derived GFP cells (green) and blood-derived TdTomato cells (orange) within the lesion boundary (A) and peri-lesion area (B). These images were generated using IMARIS software from confocal z-stack data. TdTomato cells display distinct morphological changes in the peri-lesion. They appear less round with more cellular processes when compared to GFP cells. **(C-E)** Quantification of the total volume of fluorescence (C), average sphericity (D), and average surface area (E) for GFP+ and RFP+ cells in the boundary and peri-lesion areas were determined by analyzing 3D confocal z stacks using IMARIS software. Data represented as mean± SEM (N=3-4, *p < 0.05, **p < 0.01, ***p < 0.001).

## Discussion

Neuroinflammation associated with TBI is a significant factor contributing to morbidity and mortality through complex secondary injury processes [3, 19, 20]. Traditionally, the skull has been viewed primarily as a static protective barrier for the brain. However, recent studies have transformed this perspective, identifying the skull-bone marrow as a reactive hematopoietic niche capable of contributing to and directing leukocyte trafficking into the brain [7, 21]. This emerging concept suggests that skull-bone marrow cells may be crucial in modulating brain homeostasis and neuroinflammation, particularly following injuries. While it is well-established that TBI disrupts the blood-brain barrier, leading to immune cell infiltration from the bloodstream [6, 22], the proportion and the identity of immune cells that migrate from skull bone marrow to brain parenchyma after injury remain uncertain. Using the calvarium bone-flap transplantation technique and controlled cortical impact (CCI) model of TBI, our study aimed to investigate whether calvarium-derived immune cells infiltrate the brain following TBI and differentiate them from their blood-derived counterparts. To the best of our knowledge, this study is the first to demonstrate that skull-bone marrow cells infiltrate the brain after traumatic injury, potentially influencing the pathogenesis of TBI.

Studying the effect of skull bone marrow in TBI poses significant challenges due to the intricate nature of immune cell trafficking and the diverse responses elicited by different TBI models. The controlled cortical impact (CCI) model is commonly used to study TBI, as it allows for precise control over the location and severity of the injury [6, 11, 12, 22, 23]. However, this model involves direct manipulation of the skull tissue, which can pose challenges, particularly in understanding the influence of proximal skull tissue on injury progression and recovery [21]. Despite the craniectomy involved in our model, we found that immune cells from the calvarium still infiltrate the brain following injury. This indicates that removing a 4mm craniectomy over the right parietal-temporal cortex of the skull may have a limited impact on the overall role of posterior calvarium skull bone marrow cells in the immune response. Additionally, by performing the CCI injury two weeks after a skull transplant, we allowed the transplanted bone to integrate and stabilize, minimizing additional confounding factors. Our findings suggest that, even with manipulating the skull in the CCI model, the calvarium remains a crucial source of immune cells that infiltrate the brain and contribute to the inflammatory response. The use of the CCI model thus proves effective in studying the skull-brain axis of immunity following trauma, highlighting the model’s utility in revealing the pivotal role of skull-bone marrow-derived immune cells in brain injury responses.

Recent discoveries revealed the existence of vascular channels that connect the skull bone marrow and the brain surface, enabling their direct local interaction and the migration of myeloid cells through the meninges without using systemic circulation [24]. A substantial pool of monocytes and neutrophils have been identified at the CNS borders, replenished from the skull and vertebral bone marrow. This underscores the skull bone marrow’s role in maintaining immune surveillance at these critical interfaces[7]. These studies support our findings that calvarium-derived immune cells consist of neutrophils and monocytes and that calvarium-derived macrophages exhibit distinct phenotypic responses, spatial distribution, and morphological characteristics compared to blood-derived macrophages. This indicates that the origin of infiltrating immune cells may play a significant role in regulating the cellular milieu in the damaged brain, which could help drive the neuroinflammatory response to injury.

Our findings reveal distinct phenotypic responses between blood-derived and skull-bone marrow-derived macrophages following TBI. Blood-derived macrophages, characterized by pro-inflammatory expression of CD86, were dispersed throughout the peri-lesion area, suggesting their involvement in propagating inflammation. In contrast, skull-bone marrow-derived macrophages, which predominantly exhibited an anti-inflammatory phenotype, were primarily located at the lesion boundary, indicating a potential role in the containment and resolution of the inflammation. This distinction is supported by previous studies demonstrating a unique transcriptomic profile for mouse skull bone marrow distinct from vertebral bone marrow during both homeostasis and disease conditions [8]. In addition, Cugurra et al. have reported that blood-derived monocytes infiltrating the CNS following spinal cord injury or experimental autoimmune encephalomyelitis are more pro-inflammatory than their non-blood-derived counterparts [7]. However, further studies designed to interrogate the phenotypic subsets, such as scRNAseq and/or multi-dimensional flow cytometry, are needed to confirm our results. These insights emphasize the divergent nature of calvarium- and blood-derived monocyte/macrophages following brain injury and highlight the need to explore their role in the injury and recovery response.

Efferocytosis, the process of clearing dead cells, is crucial for resolving inflammation and promoting tissue repair after TBI [6]. Our previous studies have shown an upregulation of “find-me” signal receptors (S1pr1 and Cx3cr1), engulfment receptor (MerTK), and bridging molecules (Gas6 and Pros1) in the damaged cortex, indicating active efferocytosis after trauma. Using GFP chimeric bone marrow transplants, we found that a proportion of peripheral-derived monocyte/macrophages infiltrating the brain following CCI injury contribute to the clearance of apoptotic debris[6]. In this previous study, we did not distinguish between blood-versus skull-bone marrow-derived macrophages. However, in the current study, we found GFP calvarium-derived cells express MerTK, and they participate in the clearance of apoptotic debris in the damaged cortex. These findings support our claim that this population may be involved in reducing inflammation and promoting repair. Our findings are the first to report efferocytosis by calvarium-derived monocyte/macrophages following brain injury. Future studies delineating the molecular pathways involved in efferocytosis between cells of different origins are required.

In conclusion, our findings elucidate key characteristic features of infiltrating skull-bone marrow-derived monocytes/macrophages in the cortical milieu after injury. Their ability to adopt a predominant anti-inflammatory phenotype and engage in efferocytosis suggests they may be important players in resolving inflammation and promoting tissue repair. These insights pave the way for future studies investigating their causal role in TBI outcomes.

## Acknowledgments

This work was supported by the National Institute of Neurological Disorders and Stroke of the National Institutes of Health, R01NS121103 (MHT), R01NS112541 (MHT), R01NS119540 (MHT),

## References

1. in Data Integration in Learning Health Care Systems for Traumatic Brain Injury: Proceedings of a Workshop. 2024: Washington (DC).

2. Whiting, M.D., A.I. Baranova, and R.J. Hamm, Cognitive Impairment following Traumatic Brain Injury, in Animal Models of Cognitive Impairment, E.D. Levin and J.J. Buccafusco, Editors. 2006: Boca Raton (FL).

3. Theus, M.H., Neuroinflammation and acquired traumatic CNS injury: a mini review. Front Neurol, 2024. 15: p. 1334847.

4. Alam, A., et al., Cellular infiltration in traumatic brain injury. J Neuroinflammation, 2020. 17(1): p. 328.

5. Russo, M.V. and D.B. McGavern, Inflammatory neuroprotection following traumatic brain injury. Science, 2016. 353(6301): p. 783–5.

6. Eman Soliman, J.L., Erwin Kristobal Basso, Ilana Gershenson, Jing Ju, Jatia Mills, Caroline Jager, Alexandra M. Kaloss, Mohamed Elhassanny, Daniela Pereira, Michael Chen, Xia Wang, Michelle H. Theus, Efferocytosis is restricted by axon guidance molecule EphA4 via ERK/Stat6/Mertk signaling following brain injury. researchsquare.com, 2023.

7. Cugurra, A., et al., Skull and vertebral bone marrow are myeloid cell reservoirs for the meninges and CNS parenchyma. Science, 2021. 373(6553).

8. Mazzitelli, J.A., et al., Cerebrospinal fluid regulates skull bone marrow niches via direct access through dural channels. Nat Neurosci, 2022. 25(5): p. 555–560.

9. Tian, Y., et al., The Underlying Role of the Glymphatic System and Meningeal Lymphatic Vessels in Cerebral Small Vessel Disease. Biomolecules, 2022. 12(6).

10. Truett, G.E., et al., Preparation of PCR-quality mouse genomic DNA with hot sodium hydroxide and tris (HotSHOT). Biotechniques, 2000. 29(1): p. 52, 54.

11. Soliman, E., et al., Conditional Deletion of EphA4 on Cx3cr1-Expressing Microglia Fails to Influence Histopathological Outcome and Blood Brain Barrier Disruption Following Brain Injury. Front Mol Neurosci, 2021. 14: p. 747770.

12. Kowalski, E.A., et al., Peripheral loss of EphA4 ameliorates TBI-induced neuroinflammation and tissue damage. J Neuroinflammation, 2019. 16(1): p. 210.

13. Holl, E.K., Generation of bone marrow and fetal liver chimeric mice. Methods Mol Biol, 2013. 1032: p. 315–21.

14. Okyere, B., et al., Endothelial-Specific EphA4 Negatively Regulates Native Pial Collateral Formation and Re-Perfusion following Hindlimb Ischemia. PLoS One, 2016. 11(7): p. e0159930.

15. Brickler, T., et al., Nonessential Role for the NLRP1 Inflammasome Complex in a Murine Model of Traumatic Brain Injury. Mediators Inflamm, 2016. 2016: p. 6373506.

16. Brickler, T.R., et al., Angiopoietin/Tie2 Axis Regulates the Age-at-Injury Cerebrovascular Response to Traumatic Brain Injury. J Neurosci, 2018. 38(45): p. 9618–9634.

17. Hazy, A., et al., Divergent age-dependent peripheral immune transcriptomic profile following traumatic brain injury. Sci Rep, 2019. 9(1): p. 8564.

18. Soliman, E., et al., Efferocytosis is restricted by axon guidance molecule EphA4 via ERK/Stat6/MERTK signaling following brain injury. J Neuroinflammation, 2023. 20(1): p. 256.

19. Corrigan, F., et al., Neurogenic inflammation after traumatic brain injury and its potentiation of classical inflammation. J Neuroinflammation, 2016. 13(1): p. 264.

20. Kumar, A. and D.J. Loane, Neuroinflammation after traumatic brain injury: opportunities for therapeutic intervention. Brain Behav Immun, 2012. 26(8): p. 1191–201.

21. Goodman, G.W., et al., The emerging importance of skull-brain interactions in traumatic brain injury. Front Immunol, 2024. 15: p. 1353513.

22. Kowalski, E.A., et al., Monocyte proinflammatory phenotypic control by ephrin type A receptor 4 mediates neural tissue damage. JCI Insight, 2022. 7(15).

23. Cash, A., et al., Endothelial deletion of EPH receptor A4 alters single-cell profile and Tie2/Akap12 signaling to preserve blood-brain barrier integrity. Proc Natl Acad Sci U S A, 2023. 120(41): p. e2204700120.

24. Herisson, F., et al., Direct vascular channels connect skull bone marrow and the brain surface enabling myeloid cell migration. Nat Neurosci, 2018. 21(9): p. 1209–1217.

